# Ischemia-reperfusion and acidosis induce retrotransposon derepression in the mouse brain

**DOI:** 10.64898/2026.07.31.741803

**Authors:** Youfang Zhou, Angela Barattini, Xiang-ming Zha, Wenyan Sun

## Abstract

Retrotransposons are repetitive DNA elements normally suppressed through epigenetic mechanisms. In aging and neurodegenerative diseases, abnormal retrotransposon activation occurs and leads to neurotoxicity. However, whether ischemic stroke induces retrotransposon activation remains unclear. Here, we investigated whether cerebral ischemia triggers dysregulation of retrotransposon in the brain. Since brain ischemia leads to tissue acidosis, we further examined whether acidosis contributes to this response. We performed 45 min of transient middle cerebral artery occlusion (tMCAO) followed by reperfusion on adult wild-type mice, then measured expression of retrotransposons LINE-1, IAP, ETn2 and SINE B2 in ipsilateral brain tissue using RT-qPCR. Retroviral GAG protein expression was examined by Western blotting. In vitro, we exposed neuro-2a (N2A) cells to acidic extracellular pH (6.4 or 6.0) and retrotransposon transcripts were analyzed. We found that ischemia-reperfusion increased expression of IAPEz-gag and ETn2 in ipsilateral ischemic brain tissue at 24 h, with stronger induction of LINE-1 Orf1, IAPEz-gag and B2 in infarct than peri-infarct brain regions. Western blotting analysis revealed dynamic changes in GAG proteins, with no detectable changes at 6 h of reperfusion followed by reduced levels of the precursor Pr65 and mature capsid p30 products at 24 h. In N2A cells, extracellular acidosis induced time-dependent increases in LINE-1 Orf1, IAPEz-gag and B2 transcripts. These findings identify retrotransposon dysregulation as a molecular feature of ischemic brain injury and provide a framework for further investigating its role in stroke pathology.

## Introduction

Ischemic stroke is one leading cause of the death and long-term neurological disability worldwide [1]. Reperfusion therapies greatly improve cerebral blood flow restoration but the clinical outcome in patients remains highly variable. This variability highlights the need for a better understanding of the molecular mechanism underlying ischemia-reperfusion injury, as well as identification of early therapeutic targets that could improve stroke-induced neurological recovery.

Retrotransposons are repetitive genetic elements that account for about 35 % of the human and mouse genomes [2]. Most retrotransposons have accumulated mutations and lost mobilization capacity, but a subset remains transcriptionally active and retains the ability to mobilize. For instance, the long interspersed nuclear elements-1 (LINE-1) are the only active autonomous retrotransposons in both human and mouse genomes and the retrotransposition-competent LINE-1 elements are present in both human and mouse genomes [3]. The intracisternal A-particle (IAP) is one of the most transpositional active families in mouse genomes, which contains about 4000 copies of full-length or nearly full-length IAP [4]. Normally, retrotransposons are tightly controlled by epigenetic and post-transcriptional silencing mechanisms [5]. However, they can become derepressed during cellular stress and potentially contribute to genomic instability, altered gene expression and cellular dysfunction. Increasing evidence have linked retrotransposon dysregulation to multiple neurodegenerative disorders, including Alzheimer’s disease [6, 7], Parkinson’s disease [8, 9], amyotrophic lateral sclerosis (ALS), and frontotemporal dementia (FTD) [10]. These findings suggest that retrotransposon activation may represent an important mechanism contributing to neuronal injury.

In the context of cerebral ischemia, however, the activation of retrotransposons remains poorly investigated. Previous studies have reported increase expression of LINE-1 and IAP in mouse hippocampus following global cerebral ischemia at 24 h of reperfusion [11]. The mouse endogenous retrovirus-like retrotransposon VL30 expression is upregulated after focal ischemia at 20 h of reperfusion [12]. In this study, we applied a tMCAO mouse model to investigate retrotransposon expression in post-ischemic brain tissue, including micro-dissected mixed infarct and peri-infarct regions, infarct and peri-infarct regions at 24 h post-stroke. Given that ischemic stroke is accompanied by brain tissue acidosis [13], we employed an in vitro acidosis model exposing neuro-2a (N2A) neuronal cells to extracellular different pH to determine whether ischemia-associated acidosis contributes to retrotransposon activation.

## Materials and Methods

### Experimental animals

All animal procedures were conducted in accordance with and approved by the Tulane University Animal Care and Use Committee. Adult C57/BL6 mice were housed under controlled conditions (22 ± 2 °C, 50-60 % humidity; 12 h light/dark cycle) with ad libitum access to food and water. Animals were randomly assigned to experimental groups and acclimatized prior to experimentation. At the experimental endpoint, mice were euthanized using approved humane methods, and brain tissues were rapidly dissected, frozen, and stored at -80 °C for subsequent analyses.

### Transient focal ischemia

The transient middle cerebral artery occlusion (tMCAO) was induced in adult mice using the intraluminal filament model as we described previously [14, 15]. Mice were anesthetized with 1.5 % isoflurane in a mixture of 70 % N_2_O and 30 % O_2_ and kept at a stable body temperature (37 ± 0.5 °C) throughout surgery. A silicone-coated monofilament was inserted through the external carotid artery and carefully advanced into the internal carotid artery to block the origin of the middle cerebral artery. Cerebral blood flow was monitored continuously using laser Doppler flowmetry (MoorVMS-LDF2), and occlusion was considered successful when blood flow dropped by at least 80 % from baseline. After 45 min, the filament was gently removed to retore blood flow, which was confirmed by reperfusion. Sham-operated mice underwent the same procedure without insertion of the filament. After surgery, animals were allowed to recover with free access to food and water. Mice were excluded from the analysis if they failed to maintain a stable blood flow reduction (5-20 % of baseline during occlusion) or have at least 55 % reperfusion following suture removal.

### RNA extraction, cDNA and RT-qPCR assay

Total RNA was extracted from homogenized brain samples using TRIzol and then quantified by Nanodrop. First-strand cDNA was synthesized from 500 ng of total RNA using reverse transcription kit according to the manufacturer’s instructions (Bio-rad, #1725035). qPCR was performed using SYBR Green Supermix (Bio-rad, #1725124) on Applied Biosystems QuantStudio 3 in a total reaction volume of 20 µL, containing cDNA template, primers, and SYBR Green qPCR master mix. Thermal cycling conditions were 95 °C for 30 s (initial denaturation) followed by 40 cycles of denaturation at 95 °C for 3 s and annealing/extension at 60 °C for 30 s. Relative gene expression was calculated by the 2−^ΔΔCt^ method after normalization to housekeeping genes. Results are presented as fold change relative to control groups. The primers for each target are shown in Table 1.

**Table 1:**
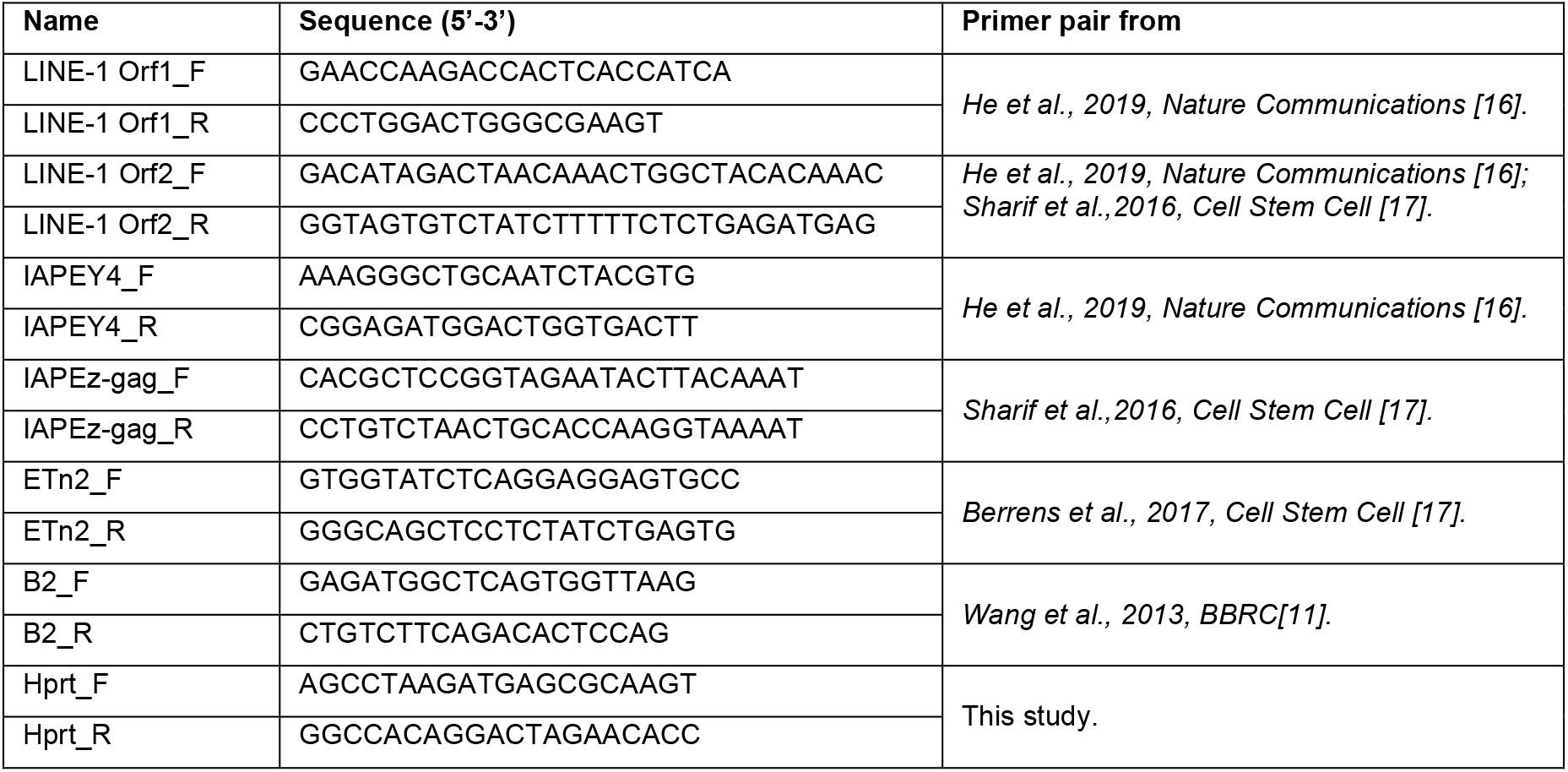
Primer sequences that used for RT-qPCR.

### Western blotting

Brain tissues were lysed in lysis buffer (1 % Triton, 0.4 % SDS, proteinase inhibitor, phosphatase inhibitor in 1 x PBS), then homogenized, sonicated and centrifuged at 16,000 x g for 10 min 4 °C. The supernatant was collected, and the protein concentration was quantified using the Bio-Rad protein assay. The denatured protein samples were separated with 12 % gels and transferred to nitrocellulose membranes. Membranes were blocked in blocking buffer (0.1 % casein, 0.2 x PBS, pH 7.2-7.6) for 30-60 min at RT, then incubated overnight at 4 °C with the following primary antibodies: anti-MLV GAG antibody (1:1000, Abcam, catalog # ab100970), anti-beta tubulin, E7 (1:10,000, Developmental Studies Hybridoma Bank). Following PBST wash, membranes were incubated with secondary antibodies for 1 h at RT (Donkey anti-rabbit IRDye 800CW, 1:10,000, Li-Cor, catalog # 926-32213; Donkey anti-mouse IRDye 680LT, 1:10,000, Li-Cor, 926-68022). After 3 times washes, the membranes were imaged using a Bio-Rad ChemiDoc MP Imaging System. The band intensities were quantified by ImageJ and normalized to loading controls.

### Statistical analysis

All data are presented as mean ± standard error of the mean (SEM) unless otherwise stated. Statistical analyses were performed using GraphPad Prism. Comparisons between two groups were assessed using an unpaired two-tailed Student’s t-test. For comparisons involving more than two groups, one-way analysis of variance (ANOVA) followed by Tukey’s post hoc multiple comparisons test was applied. Statistical significance was defined as p < 0.05.

## Result

### Retrotransposon transcripts are elevated in ipsilateral brain tissue 24 h after reperfusion

To determine whether acute ischemia induces retrotransposon activation, we performed 45 min tMCAO on 3-4 months old adult male wild type mice, collected ipsilateral and contralateral brain tissues at 24 h after reperfusion, and quantified retrotransposon transcript levels by RT-qPCR. In this study, we focused on retrotransposon families previously implicated in neurotoxicity and neurodegeneration: IAP, LINE-1, and early transposon (ETn) elements. Because tMCAO predominantly affects the striatum and primary sensorimotor cortex, we pooled these ipsilateral brain regions including peri-infarct and infarct regions for analysis.

LINE-1 contains two open reading frames, LINE-1 ORF1 and ORF2, within its full-length transcript. ORF1 encodes an RNA-binding protein, whereas ORF2 encodes endonuclease and reverse transcriptase activities and both ORF1p and ORF2p are essential for retro-transposition [18]. IAPEz is an evolutionarily young and active IAP subfamily. Its derived gag sequence is a highly conserved region shared across multiple IAP elements and provide a broad readout of IAP transcriptional activation [19]. RT-qPCR analysis showed significantly increased expression of IAPEz-gag and ETn2 transcripts in ipsilateral brain tissue from tMCAO mice relative to sham controls, whereas LINE-1 Orf1 transcripts showed an upward trend (Fig. 1A). We also compared basal retrotransposon expression in young (3-4 months) and aged (17-19 months) wild type male mice without surgery. Consistent with previous reports demonstrating age-associated retrotransposon derepression [20], IAPEz-gag, LINE-1 Orf1 and ETn2 transcript levels were highly elevated in aged brains compared with young controls (Fig. 1B). These data suggest that ischemia-reperfusion injury is associated with dysregulated retrotransposon transcripts in the post-ischemic brain.

**Figure 1.**
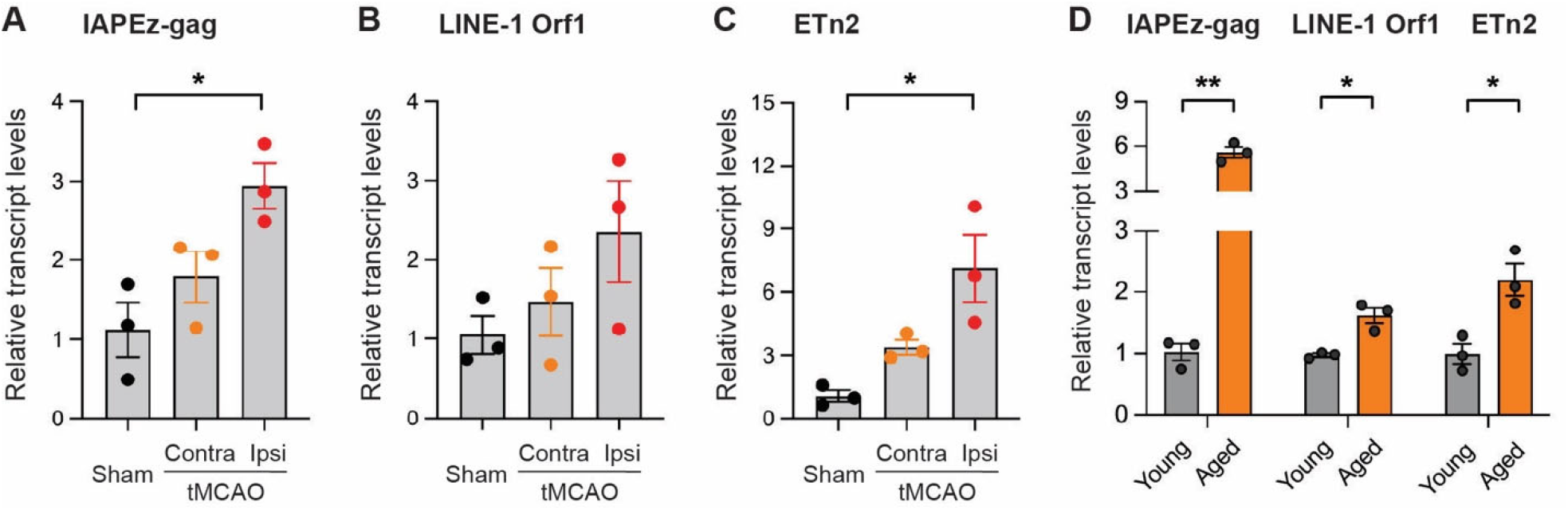
Increased retrotransposon expression in ipsilateral brain tissue following acute ischemic stroke. Relative expression of retrotransposons in mixed peri-infarct and infarct brain tissue was measured by RT-qPCR. **(A)** IAPEz-gag; **(B)** LINE-1 Orf1; **(C)** ETn2. **(D)** Comparison of retrotransposon expression in aged (17-19 months) versus young (3-4 months) wild-type (WT) mice. Gene expression levels were normalized to Hprt and calculated using the 2^^−ΔΔCt^ method. Data are shown as mean ± SEM. \**p* < 0.05; \*\**p* < 0.01. Contra, contralateral region; Ipsi, ipsilateral region. n = 3 male mice per group.

### Retrotransposons are derepressed in infarct regions following reperfusion

In ischemic stroke, the infarct region tissue has been irreversibly injury due to prolonged reduction in cerebral blood flow and oxygen deprivation. Surrounding this region is the peri-infarct region/penumbra zone, which consists of hypoperfused but potentially viable tissue. This distinction is clinically relevant because the aim of acute therapeutic interventions is to preserve peri-infarct tissue and limit infarct expansion [21, 22]. To determine whether retrotransposon expression differs between infarct and peri-infarct regions following ischemia-reperfusion injury, we performed 45 min tMCAO on WT male mice. At 24 h after reperfusion, ipsilateral peri-infarct and infarct, and corresponding region matched contralateral tissues were micro-dissected separately. Total RNA was extracted, and quantitative RT-PCR was performed using primers targeting LINE-1, IAP, B2 and ETn2 elements. To further investigate IAP activation, we quantified IAPEY4 which is a relatively young and transcriptionally active subfamily member.

LINE-1 Orf1 and Orf2 transcripts exhibited modest increase in peri-infarct region relative to sham and peri-infarct-matched contralateral regions. In contrast, LINE-1 Orf1 transcripts were significantly elevated in infarct tissue compared with sham and contralateral infarct-matched regions (Fig. 2A) and LINE-1 Orf2 shows modest increased in infarct regions (Fig. 2B). Similarly, IAPEz-gag and IAPEY4 transcripts showed intermediate increases in peri-infarct tissue and significant upregulation within the infarct core (Fig. 2C-D). SINE B2 shows significant increase in infarct region compared to sham controls and peri-infarct region (Fig. 2E), whereas ETn2 transcripts were not significantly changed in either region (Fig. 2F). These findings demonstrated that ischemia-reperfusion injury is associated with increased expression of LINE-1 Orf1, IAPEz-gag and B2 elements in infarcted brain tissue at 24 h. The intermediated induction observed in peri-infarct regions suggest that retrotransposon dysregulation may occur early during ischemic stress.

**Figure 2.**
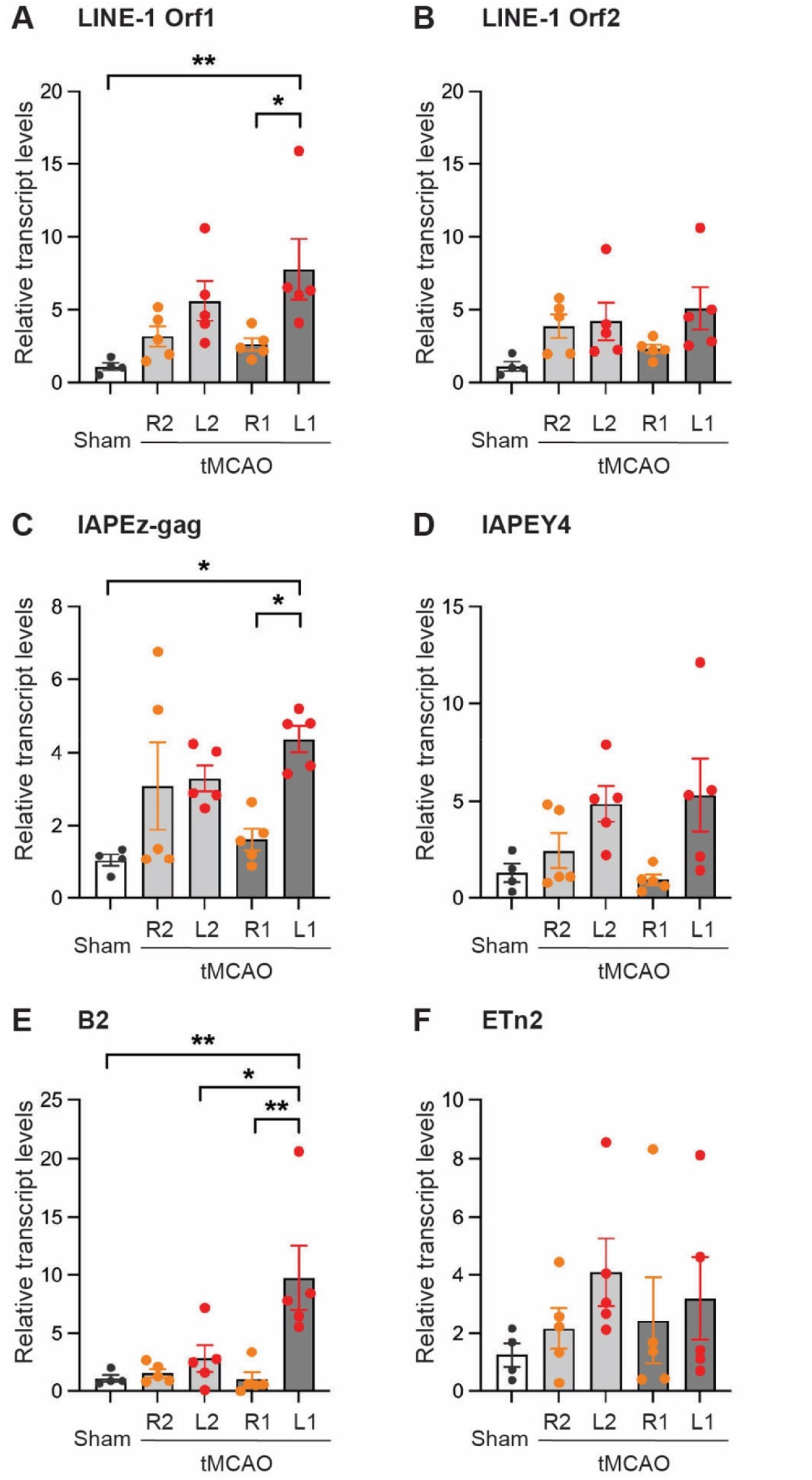
Regional specific retrotransposon expression in peri-infarct and infarct regions following ischemic stroke. Relative expression of retrotransposons in peri-infarct and infarct brain tissue was analyzed by RT-qPCR. **(A)** LINE-1 Orf1; **(B)** LINE-1 Orf2; **(C)** IAPEz-gag; **(D)** IAPEY4; **(E)** B2; **(F)** ETn2. Gene expression levels were normalized to Hprt and calculated using the 2^^−ΔΔCt^ method. Data are shown as mean ± SEM. \**p* < 0.05; \*\**p* < 0.01. R2, contralateral region corresponding to the peri-infarct area; L2, ipsilateral peri-infarct region; R1, contralateral region corresponding to the infarct region; L1, ipsilateral infarct region. n = 4-5 male mice per group, 3-4 months of age.

### Dynamic changes in retroviral GAG-immunoreactive proteins following ischemic brain injury

Endogenous retroviruses (ERVs) are the integrated proviral form of ancestral exogenous retroviruses in germline cells. As a result, many ERVs still have retroviral-like coding sequences, including gag, pol and env domains [23]. In exogenous retroviruses such as murine leukemia virus (MLV), the gag open reading frame encodes a precursor polyprotein (Pr65) that is proteolytically cleaved to matrix (MA p15), capsid (CA, p30) and nucleocapsid protein (NC, p10) and p12 proteins [24]. To determine whether retroelement-associated GAG proteins are altered after ischemic injury, we performed tMCAO in mice, isolated ipsilateral (Ipsi) and contralateral (Contra) brain tissues with sham control and performed western blot analysis using a commercial anti-MLV GAG antibody.

At 6 h reperfusion, ipsilateral ischemic tissue showed no significant changes in either the precursor GAG protein Pr65 or its cleavage products compared with contralateral or sham brain tissues (Fig. 3A-B). There’s also no accumulation of lower-molecular-weight GAG-reactive bands p30 protein in either ipsilateral or contralateral tissue at this time point (Fig. 3A). At 24 h reperfusion, the Pr65 was significantly reduced in ipsilateral tissue. In addition, p30 protein was significantly increased in contralateral tissue but decreased in the ipsilateral tissue, whereas the p15-reactive band remain unchanged (Fig. 3C-E). These findings suggest that GAG-immunoreactive proteins undergo dynamic changes during ischemic injury, characterized by an early period with no detectable changes followed by a later reduction in both precursor protein Pr65 and mature capsid p30.

**Figure 3.**
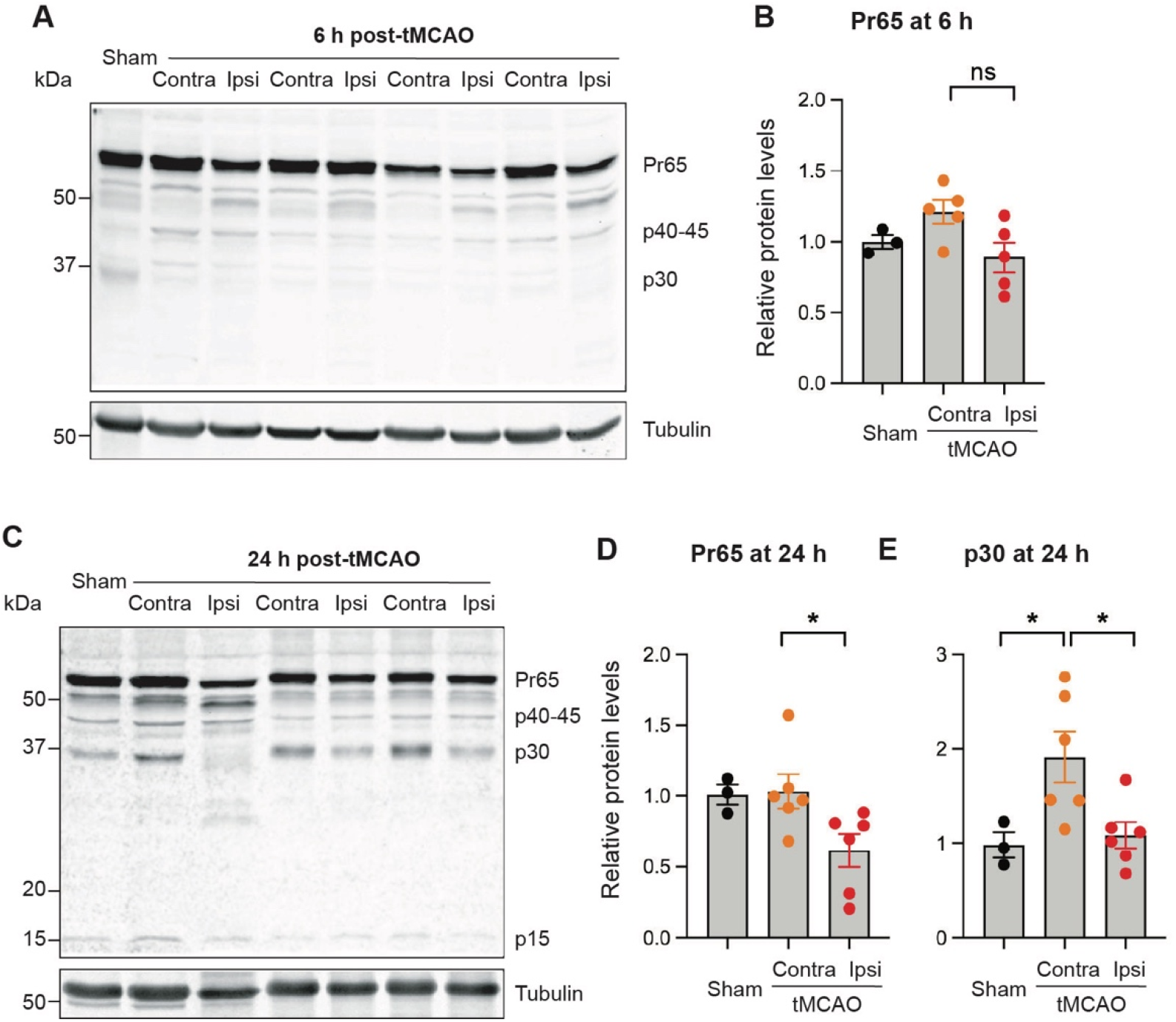
Dynamic changes of retroviral GAG-immunoreactive proteins following acute ischemic brain injury by Western blotting analysis. **(A-B)** At 6 h post-reperfusion, no changes in cleaved products. **(C-E)** At 24 h post-reperfusion, decreased levels of precursor Pr65 and decreased mature capsid (p30) were observed in ischemic brain tissue. Quantification of western blot signals: **(B)** Pr65 at 6 h, **(D)** Pr65 at 24 h, **(E)** p30 at 24 h. Protein levels were normalized to tubulin. Contra, contralateral region; Ipsi, ipsilateral region. Data are shown as mean ± SEM. \**p* < 0.05. n = 3-6 male mice per group, 3-4 months of age.

### Acidosis induces LINE-1 Orf1, IAPEz-gag, and B2 transcript accumulation in neuronal cells

Cerebral ischemic stroke impairs oxygen delivery to brain tissue, leading to a shift metabolism with accumulation of lactic acid and elevated tissue CO_2_ levels, resulting in tissue acidosis. This ischemic acidosis contributes significantly to neuronal dysfunction, cellular injury, and subsequent neurological deficits [25]. To determine whether acidosis induces aberrant expression of retrotransposon in neurons, we treated mouse neuroblasts Neuro-2a (N2A) cells with physiological pH (7.4, control) or acidic conditions (pH 6.4 and 6.0). To assess temporal changes in retrotransposon transcription, we harvested cells for 6 h and 24 h following pH treatment, extracted RNA and analyzed by qRT-PCR to quantify transcripts of LINE-1 Orf1, LINE-1 Orf2, IAPEz-gag, IAPEY4, B2 and ETn2 elements.

We found that neuronal acidic stress induced the upregulation of several retrotransposon transcripts. After 6 h of exposure to pH 6.4 or 6.0, LINE-1 Orf1 and Orf2 expression remained unchanged (Fig. 4A). IAPEz-gag showed an increasing trend but IAPEY4 was decreased (Fig. 4B). B2 transcript levels was significantly increased (Fig. 4C), and ETn2 was reduced at pH 6.4 (Fig. 4D). At 24 h, LINE-1 Orf1 was significantly upregulated under both acidic conditions, whereas LINE1-Orf2 persisted unchanged (Fig. 4E). IAPEz-gag transcripts were significantly increased under pH 6.0, whereas IAPEY4 expression was not changed (Fig. 4F). B2 transcripts exhibited a further marked increases (Fig. 4G). Etn2 transcript levels were not affected (Fig. 4H). These results indicate that neuronal acidosis selectively activates retrotransposon subfamilies, particularly LINE-1 Orf1, IAPEz-gag and B2 in a time-dependent manner.

**Figure 4.**
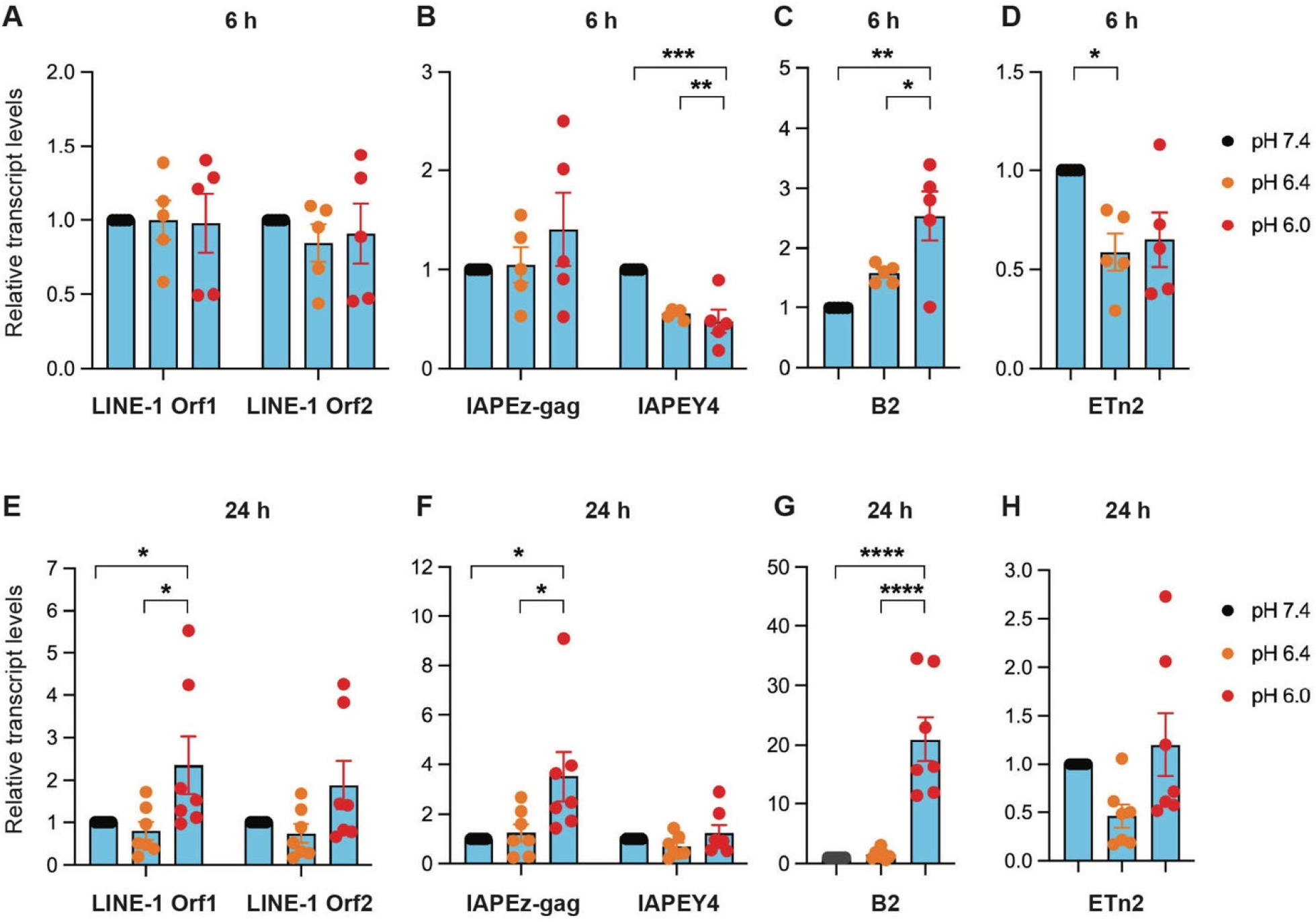
Acidic extracellular pH induces retrotransposon expression in N2A cells. N2A cells were exposed to pH 7.4 (control), 6.4, or 6.0 for 6 h or 24 h followed by RT-qPCR analysis. Relative expression levels at 6 h of **(A)** LINE-1 Orf1 and Orf2; **(B)** IAPEz-gag and IAPEY4; **(C)** B2; **(D)** ETn2. Relative expression levels at 24 h of **(E)** LINE-1 Orf1 and Orf2; **(F)** IAPEz-gag and IAPEY4; **(G)** B2; **(H)** ETn2. Gene expression levels were normalized to Hprt and are presented as fold change using the 2^^−ΔΔCt^ method. Data are presented as mean ± SEM. \**p* < 0.05; \*\**p* < 0.01, \*\*\**p* < 0.001, \*\*\*\**p* < 0.0001. n = 5-7 independent experiments.

## Discussion

### Retrotransposon derepression is enriched within infarct tissues following ischemia-reperfusion injury

Our study identifies retrotransposon derepression as a molecular feature of transient cerebral ischemia-reperfusion injury. The elevated expression of LINE-1 Orf1, IAPEz-gag and B2 elements in young adult tMCAO mice suggest that ischemic injury disrupts mechanisms for transposable element silencing in the brain. The comparison between young and aged brains further strengthens the relevance of these findings. Aging is well established to promote retrotransposon activation through epigenetic dysregulation and is a major risk factor for ischemic stroke [26, 27]. The elevated LINE-1 Orf1, IAPEz-gag, and ETn2 expression we observed in aged mouse brain is consistent with transposable element dysregulation associated with aging and neurodegeneration.

Our findings further demonstrate that activation of retrotransposon exhibit region differences within the ipsilateral brain, with the strongest induction in the infarct core and a modest increase detected in peri-infarct tissue. The significant upregulation of LINE-1 Orf1, IAPEz-gag, and B2 elements in infarcted regions suggests that severity of ischemia is associated with widespread dysregulation of retroelements. The mild increase we observed in peri-infarct tissue is interesting, as this region contains metabolically stressed but potentially salvageable cells. This expression pattern provides the possibility that retrotransposon activation occurs early during ischemic injury progression and may contribute to inflammatory signaling or genomic instability before irreversible tissue damage develops. ETn2 transcripts were not significantly changed following ischemia-reperfusion injury, suggesting differential regulation among retrotransposon families. This selective response may reflect distinct transcription factor dependencies, or differences in temporal activation and cell-type specificity. Because our analyses were performed on bulk brain tissue, the cellular sources of retrotransposon induction remain unresolved, particularly given the heterogeneous responses of neurons, glia, and infiltrating immune cells after stroke. Single-cell transcriptomic approaches will help to define cell-type-specific retrotransposon dynamics following ischemic injury. The other limitation of our study is that all young adult mice were male. Given the well-established sex differences in both experimental models and clinical populations [28], it remains unclear whether the profile of retrotransposon dysregulation differs in females. Inclusion of both sexes in future studies will be important for evaluating the generalizability of these findings.

### Alterations of retroviral GAG-immunoreactive proteins during ischemic brain injury

We observed dynamic changes in GAG-immunoreactive proteins following ischemia-reperfusion, characterized by no detectable changes at 6 h following reperfusion and reductions in both precursor Pr65 and mature capsid p30 products at 24 h. Their reduction likely reflects decreased abundance of GAG-expressing cells and/or reduced protein expression within injured tissue. Interestingly, the capsid p30 was elevated in contralateral tissue compared with sham controls, suggesting that ischemic injury may induce retroviral-like protein responses not only within the infarcted regions but also in contralateral brain regions. ERVs share the same genomic organization as exogenous retroviruses, including gag, pol, and env genes [29]. ERVs can retain expression capacity, providing a molecular basis for antigenic similarity and potential antibody cross-reactivity [24]. The anti-MLV GAG antibody used was originally developed against murine leukemia virus proteins and likely recognizes conserved retroviral epitopes shared among multiple endogenous retroelement families. Therefore, although the detected bands migrated at molecular weights consistent with retroviral GAG proteins, we cannot conclusively assign them to a specific endogenous retrovirus or retrotransposon family. Multiple murine ERV and LTR retrotransposon families, including IAPs, endogenous MLV-related elements, ETn/MusD elements, and ERVK-related sequences, retain conserved retroviral gag domains [29, 30]. All of these may contribute to the observed immunoreactivity. Despite this technical caveat, the result showed that ischemia-reperfusion induced dynamic changes of GAG protein processing.

### Extracellular acidosis selectively induces retrotransposon expression in neuronal cells

Ischemic stroke induces brain tissue acidosis through impaired oxidative metabolism, lactate accumulation, and elevated CO_2_ levels [13]. Therefore, extracellular acidosis is a major component of the ischemic microenvironment and acidotoxicity contributes to neuronal injury. Brain acidosis during ischemia disrupts multiple homeostatic pathways, including mitochondrial function and oxidative metabolism [31], calcium and pH homeostasis [32], and chromatin organization [33]. These processes are closely linked to epigenetic stability and transposable element silencing providing several potential mechanisms through which acidosis may affect retrotransposon expression. The upregulation of LINE-1 Orf1, IAPEz-gag, and B2 under acidic conditions in N2A cells suggests that ischemia-associated acidosis contributes to retrotransposon dysregulation. However, LINE-1 Orf2, IAPEY4 and Etn2 transcripts remained unchanged at 24 h. This finding suggests that acidosis stress does not induce a global upregulation of all retrotransposons. Instead, acidosis selectively affects a subset of retrotransposons.

The selective induction of specific retrotransposon expression in this study may reflect differential sensitivity of individual retrotransposon families to acidosis and its downstream effects on transcriptional regulation. The regulatory mechanisms underlying this differential regulation warrant further investigation. We recognize that extracellular acidosis represents only one component of ischemic injury. Therefore, the *in vitro* acidosis model we used in this study captures only a subset of cellular stress responses that occur during cerebral ischemia. Combining other ischemia-relevant factors, such as hypoxia or oxygen-glucose deprivation, will provide a more comprehensive understanding of the mechanisms underlying retrotransposon dysregulation after stroke.

## Conclusion

In summary, we demonstrated that acute ischemia-reperfusion leads to retrotransposon derepression in the post-ischemic mouse brain, and that extracellular acidosis is sufficient to upregulate selective retrotransposon expression in neuronal cells. This result suggests that transient ischemia, in part through acidosis, disrupts retrotransposon silencing mechanisms. Further studies are needed to determine the mechanism by which ischemia or acid signaling leads to retrotransposon activation, and whether retrotransposon dysregulation directly contributes to neuronal injury, neuroinflammation, and functional recovery after stroke.

## Author Contributions

Y.Z. and W.S. performed mouse surgeries and conducted RT-qPCR analyses. A.B. performed Western blotting. X.Z and W.S. conceived the study, analyzed data and wrote the manuscript.

## Declarations

The authors declare no completing interests.

